# Adaptation to warm environments with a fast pace of life in a marine predatory snail

**DOI:** 10.1101/2025.06.20.660751

**Authors:** Christopher Dwane, Lisa M. Komoroske, Allison L. Rugila, Blair P. Bentley, Emma Rawson, Elizabeth Clark, Gillian Nichols, Mikayla Newbrey, Emily Bucari, Chance Yan, Jordanna Barley, Ryan Horrigan, Liam McCarthy, Nicholas Duncan, Ana Beatriz Juarez Stucker, Andrew R. Villeneuve, Brian S. Cheng

**Affiliations:** Department of Environmental Conservation, University of Massachusetts Amherst, Amherst, Massachusetts 01003, USA; Marine Biology and Ecology Research Centre, School of Biological and Marine Sciences, University of Plymouth, Plymouth PL4 8AA, UK; Department of Biological Sciences, Smith College, Northampton, Massachusetts 01063, USA; Mount Holyoke College, South Hadley, Massachusetts 01075, USA; Department of Biological Sciences, University of New Hampshire, Durham, New Hampshire 03824, USA; Department of Biology, Monmouth University, West Long Branch, New Jersey 07764, USA; Department of Marine and Environmental Sciences, Northeastern University, Boston, Massachusetts 02115, USA; Harvard Graduate School of Education, Harvard University, Cambridge, Massachusetts 02138, USA

## Abstract

Understanding how latitudinal temperature variation shapes local adaptation of life history strategies is crucial for predicting future responses to warming. Contrasting predictive frameworks explain how growth and other life history traits may respond to differing selective pressures across latitude. However, these frameworks have rarely been explored within the context of fluctuating environmental temperatures across longer (i.e., seasonal) time scales experienced in nature. Furthermore, consequences of growth differences for other aspects of fitness, including reproductive output, remain unclear. Here, we conducted a long-term (17-month) simulated reciprocal transplant experiment to examine local adaptation in two populations of the predatory marine snail *Urosalpinx cinerea* separated by 8.6° latitude (1000 km). We reared F1 offspring under two seasonally fluctuating temperature regimes (“warm” and “cold”, simulating field thermal conditions experienced by low and high latitude populations, respectively), quantifying temporal patterns in growth, maturation, and reproductive output. We identified striking divergence in life-history strategies between populations in the warm regime, with offspring from the low latitude population achieving greater growth in their first year, and high reproductive output coupled with reduced growth in their second year. In contrast, the high latitude population grew slower in their first year, but eventually attained larger sizes in their second year, at the expense of reduced reproductive output. Responses were consistent with this in the cold regime, although growth and reproductive output was reduced in both populations. Our data provides support for adaptive divergence across latitude consistent with the Pace-of-Life hypothesis, with the low latitude population selected for a fast-paced life characterized by rapid development and early reproduction. In contrast, the high latitude population exhibited slower growth and delayed maturation. Our results highlight the potential limitations of short-term comparisons of growth without considering processes over longer time scales that may exhibit seasonal temperature variation and ontogenetic shifts in energy allocation and imply a radical reshaping of physiological performance and life history traits across populations under climate change.

## Introduction

Understanding how environmental temperature shapes the evolution of life history strategies is critical to predicting how populations will be affected by a changing climate (Angilletta et al. 2002, Angilletta 2009, Lovell et al. 2023). Climatic variation across latitude represents one of the most pervasive environmental gradients on Earth, affecting all aspects of physiology and ecology (De Frenne et al. 2013). In addition to reduced mean temperatures, high latitude locations are characterised by shorter season lengths which impose constraints on growth and reproduction (Conover and Present 1990, Clarke 1993, Hochachka and Somero 2002, Varpe 2017). Low latitude locations are characterised by longer season lengths and potentially stronger intraspecific competition and predation risk (Freestone et al. 2021), which may select for altered life history strategies (Ricklefs and Wikelski 2002, Schemske et al. 2009, Varpe 2017). Latitudinal temperature variation is particularly relevant in marine ecosystems because the high heat capacity of water and relative lack of microclimate variation creates a challenging environment for aquatic ectotherms to adjust their body temperatures via behavioural thermoregulation (Gunderson and Stillman 2015, Pinsky et al. 2019, Sasaki et al. 2022). As many marine species possess broad geographic distributions, often spanning highly varied thermal conditions, it is imperative to understand how selective pressures shape the evolution of life history strategies to overcome these challenges (Deutsch et al. 2008, Conover et al. 2009, Bozinovic et al. 2011, Sanford and Kelly 2011). This information is crucial to enhance our predictions of future performance shifts under a rapidly changing climate (Deutsch et al. 2008, Lovell et al. 2023), especially as higher latitude ecosystems increasingly come to resemble those at lower latitudes (Vergés et al. 2019).

Once thought to be rare in the oceans (Sanford and Kelly 2011), local adaptation is now known to be widespread in marine species (Conover and Present 1990, Campos et al. 2009, King et al. 2018, Sasaki et al. 2022, Beaty et al. 2023). However, a mechanistic understanding of how selective pressures shape the evolution of life history strategies as a whole, as opposed to individual traits, remains elusive, in part because studies frequently consider growth rate in isolation of other traits (Metcalfe and Monaghan 2003, Biro et al. 2006, Sanford and Kelly 2011). As a result, trade-offs between growth and other traits such as reproductive output (Arendt 2011) and longevity (Metcalfe and Monaghan 2003) remain relatively poorly studied, although evidence from short-lived species suggests that large size confers greater fecundity benefits at low temperatures (Mccabe and Partridge 1997, Weetman and Atkinson 2004, Arendt 2015). Furthermore, much of our understanding of local adaptation of life history strategies is based on comparisons under constant temperatures for relatively short periods of time (i.e. common garden experiments, (Conover et al. 2009), often focusing on species possessing short generation times which may be under selection for fast growth (Yamahira and Conover 2002) or experience balancing selection across seasons (Mérot et al. 2020). Conversely, seasonality variation across latitude may play a greater selective role in longer-lived species which can postpone short term reproductive output in favour of continued growth and future reproductive success (Varpe 2017). To comprehensively understand patterns of life-history adaptation across latitude in such organisms, it is necessary to incorporate environmentally realistic temperature regimes (e.g., seasonal fluctuations) across longer time scales (e.g., months to years), and to assess multiple life history traits in tandem.

Here, we consider three contrasting theoretical models for understanding how local adaptation may drive variation in growth and corresponding reproductive traits across latitude, serving as predictive frameworks for characterising life-history strategies across populations (Figure 1). The first, which we refer to here as the Local Advantage hypothesis (Fig. 1A-C), follows directly from the adaptive principle that populations should possess traits which optimise performance under local conditions relative to other populations (i.e. a local vs. foreign comparison, *sensu* Kawecki and Ebert 2004; Yamahira and Conover 2002, Conover et al. 2009, Lovell et al. 2023). Accordingly, warm-adapted populations should possess higher growth, and consequently higher reproductive output, compared to cold-adapted populations when compared under a warm thermal regime (Fig. 1A), and vice versa under a cold thermal regime (Fig. 1B). However, thermodynamic constraints would still result in higher growth under a warm regime regardless of source population (Fig. 1C; Kawecki and Ebert 2004). A second framework, Countergradient Variation (CnGV), predicts an opposite directionality of responses (Levins 1968, Conover and Schultz 1995, Conover et al. 2009). Here, high latitude environments select for fast growth because the growing season is temporally constrained, resulting in higher growth rates in cold-adapted compared to warm-adapted populations even when compared under non-local conditions (Fig. 1D-F). CnGV, which is widely supported empirically from common garden experiments (Conover and Present 1990, Conover et al. 2009), thus represents a scenario in which selection counteracts the latitudinal effect of temperature on growth rates, allowing cold-adapted populations to grow rapidly in the short growing season and facilitate higher reproductive output (1D-F). A third predictive framework (Fig. 1G-I) may be derived from the Pace-of-Life Syndrome (POLS) hypothesis, describing the tendency for physiological, behavioural and life-history traits to positively covary along a “fast-slow” life history axis (Dammhahn et al. 2018). Fast-paced phenotypes are characterised by rapid growth and early reproduction, favoured in resource-rich, high predation environments where early reproduction increases fitness. By contrast, slow-paced phenotypes are characterised by slow but sustained growth and delayed reproductive investment, favoured in resource – poor environments where long-term survival is advantageous (Réale et al. 2010, Montiglio et al. 2018, Hämäläinen et al. 2021). Intriguingly, temperature variation across latitude covaries with several environmental variables associated with the POLS (Segev et al. 2017, Debecker and Stoks 2019, Tüzün and Stoks 2022). For instance, at low latitudes, higher predation risk and resource availability are coupled with elevated habitat temperatures, potentially facilitating higher metabolic rates and a fast-paced strategy in ectotherms, while at high latitudes, reduced temperatures may be coupled with low or seasonal resource availability, potentially promoting slow-paced phenotypes which accrue fitness benefits from late reproduction (Angilletta et al. 2004, Arendt 2011, 2015, Segev et al. 2017). Accordingly, under the POLS, correlational selection may be predicted to reinforce these divergent life-history responses across latitude, leading to preserved differences even when populations are compared under the same conditions (Fig. 1G-I).

**Figure 1:**
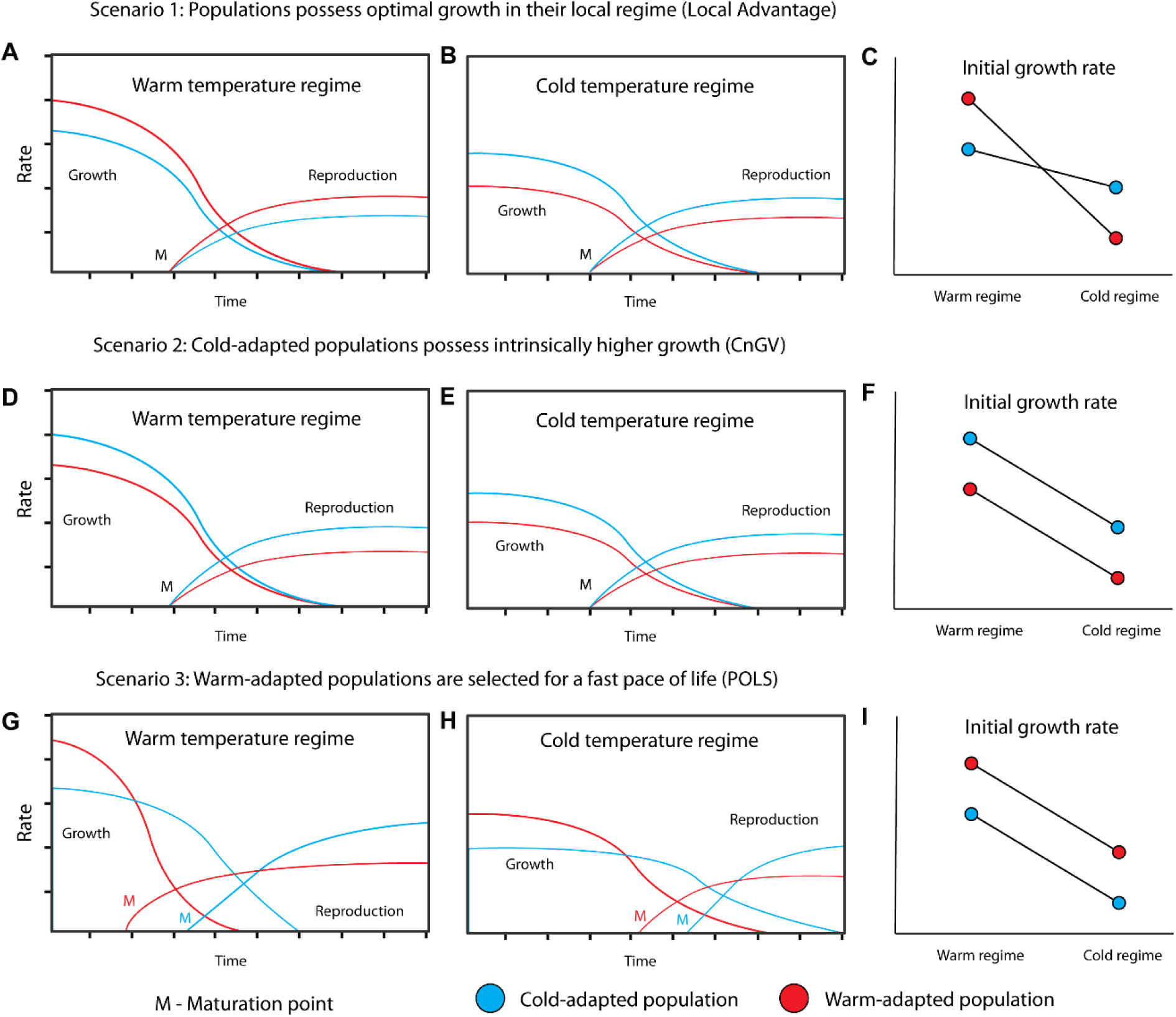
Conceptual figure outlining potential life-history responses of cold and warm- adapted populations under reciprocal transplantation between their respective thermal regimes. (A-C) Under the Local advantage hypothesis, populations tested within their local environment (red lines in A, blue lines in B) always possess optimal growth and therefore reproductive output relative to non-local populations (C). (D-F) Under Countergradient Variation, cold-adapted populations possess intrinsically higher growth rates to counteract reduced growing seasons, leading to higher growth in both warm (D) and cold (E) temperature regimes (F). This results in greater size at reproduction and thus higher reproductive output in both regimes. (G-I) Under the Pace of Life Syndrome (POLS) hypothesis, covariation of environmental temperature with resource availability, competition and predation across latitude lead to divergent selection across a “fast-slow” life-history axis, leading to warm-adapted populations which grow faster and prioritise early reproduction relative to cold-adapted populations (G-H). Cold-adapted populations initially grow more slowly (I) but reach a larger size at maturation, eventually leading to higher reproduction later in life. In all three scenarios, an underlying effect of temperature on growth rate is assumed, meaning that populations kept in the colder environment grow more slowly than those kept in the warm environment, irrespective of genetic effects (Panels C, F, I).

In this study, we explore intraspecific divergence of life-history strategies across latitude using a widespread marine predatory snail *Urosalpinx cinerea* (hereafter “*Urosalpinx*”). This species, which inhabits both the subtidal and lower intertidal zones, spans the North American Atlantic coastline from Nova Scotia to Florida (Carriker 1955) and persists across one of the strongest marine latitudinal thermal gradients in the world (Baumann and Doherty 2013). Unlike many marine taxa, *Urosalpinx* has direct development with crawl-away larvae that emerge from benthic egg capsules. Such limited dispersal capability results in low population connectivity and strong genetic structure across its native range (Bentley et al. 2024), making it an excellent candidate to study local thermal adaptation across latitude. Prior work in this system has revealed population differentiation in growth rates consistent with countergradient variation, although these comparisons were limited to shorter term common garden experiments in juveniles (Villeneuve et al. 2021a). Building upon this framework, we quantified a suite of growth and reproductive traits in populations of *Urosalpinx* using a long-term laboratory reciprocal transplant experiment. We aimed to understand interpopulation variation in life history strategies within the context of the Local Advantage, CnGV, and POLS frameworks (Fig. 1), enabling us to identify underlying adaptive mechanisms shaping local adaptation of thermal strategies across latitude.

## Methods

### 1. Broodstock collection

We collected native range adult *Urosalpinx* broodstock from a high latitude site in Great Bay, New Hampshire (43°05’29”N 70°51’55”W, henceforth NH) and a low latitude site in Beaufort, North Carolina (34°43’5”N 76°40’16”W, henceforth NC). Snails from an additional low-latitude population in Skidaway, Georgia (31°59’24”N 81°01’16”W, henceforth GA) were also collected, but we were only able to obtain offspring from a single broodstock female. Although we used these offspring in our experiment described below, we chose to exclude this population from our formal analysis due to small sample size (see Appendix S1: Table S1) and lack of replication of maternal lines. However, data from GA individuals was broadly consistent with NC snails and graphical outputs including the GA population are presented in Appendix S2: Figures S1-S2. Broodstock were collected from all sites in late April - early May 2022 and brought to the laboratory within three days of collection. We then sexed and paired broodstock and placed them into separate plastic jars inside a recirculating seawater table at 18°C ±1°C. This 275 L table, and other tables described henceforth, contained artificial seawater made using reverse osmosis deionized water and sea salt (Instant Ocean; 30 ± 1 PSU), Life support was provided with biological and mechanical filter media, foam fractionators, and temperature control (described below). Egg capsules were obtained from six females from each population and placed into tea strainers (Perma Brew tea strainers, Upton Tea Imports, Hopkinton, MA, USA) to develop at 22°C prior to placement in the reciprocal transplant experiment.

### 2. Reciprocal transplant experiment

For our reciprocal transplant design, we used two separate seawater tables, one simulating field temperatures at the New Hampshire site (henceforth referred to as “cold regime”) and the other, the North Carolina site (henceforth “warm regime”). To obtain reference temperatures for the simulations, we obtained *in-situ* data from buoys located nearby to the two sample locations from the NOAA National Data Buoy Center (further details in Appendix S1: Section S1) and generated average daily temperatures for the years 2019 – 2021. Temperature control in each seawater table was provided using a combination of chiller units (Delta Star, AquaLogic Inc., Monroe, NC, USA) and aquarium heaters connected to a thermostat, which was adjusted daily to reflect reference temperatures. This maintained temperatures within ± 1°C of set temperatures, although deviations did occur; additional information on temperature maintenance is provided in Appendix S1: Section S1.

Within ten days of hatching (on 2022-08-02), F1 juveniles from each clutch were separated into individual tea strainers and divided between the two temperature regimes. Initial temperatures at the time of addition were 22°C in both regimes, which matched reference temperatures for the cold regime at that time; temperatures in the warm regime were ramped upwards to match their respective reference temperatures (27°C) over the following five days. For both the NC and NH populations, 80 individuals from six separate maternal lines were divided between the two regimes, resulting in an initial sample size of 40 individuals per population / thermal regime combination. Due to mortality within the first three weeks following transfer (19/80 snails in the warm regime, 28/80 snails in the cold regime), snails that died within this period were replaced with individuals from the same brood. In our experience, such early mortality is typical for young *Urosalpinx* and diminished to lower levels after the first 3 weeks of the experiment. Mortality information for the remainder of the experiment is given in Appendix S1: Table S1. Data from snails which died at any stage through the experiment were excluded from further analysis. We maintained the snails on a diet of juvenile oyster (*Crassostrea virginica*, Ward Oyster, VA, USA) throughout the experiment. Fresh oysters were added to the snail enclosures on either a weekly or fortnightly basis to ensure *ad libitum* feeding, based on observed temperature dependent consumption which varied throughout the experiment. More information on animal care is given in Appendix S1: Section S2.

Beginning in March 2023, at approximately 7.5 months of age, snails were paired to allow for mating and reproduction. This date was chosen as field records indicate that it corresponds with the start of the breeding season in North Carolina (our “warm regime”), and because most snails in both regimes had achieved a sufficient size (>10mm) by this date to be sexed (Carriker 1955, Hargis 1957). For the NC and NH populations, females were outcrossed with experimental males from a different maternal line of the same population. Paired snails were housed inside larger enclosures constructed from clear plastic containers (250ml), with windows cut into the sides and covered with mesh to facilitate flow-through of seawater. More detail on pairing is given in Appendix S1: Section S3.

### 3. Size and growth rate measurements

To assess changes in size and growth rate, we measured shell length (spire tip to siphonal canal tip) throughout the experiment. The first set of measurements were obtained by photographing snails using a stereomicroscope (Leica S9i, Leica Microsystems GmbH, Wetzlar, Germany). and measured digitally using Fiji (Schindelin et al. 2012) to avoid handling damage at this small size. For subsequent measurements, taken at regular (4-6 week) intervals, we used digital callipers. Specific growth rates (SGR) between each measurement timepoint were obtained from shell lengths using the formula:

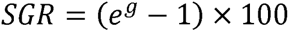

where:

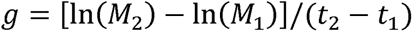

And where *M_2_* and *M_1_* represent the shell length of each snail at the current and previous timepoints (*t_2_* and *t_1_*), respectively (Jobling 1983).

We also collected tissue weight measurements using the “buoyant weight” technique (Palmer 1982) from a consistent subset of 12 individuals from each temperature regime and population combination (total n = 48 snails). At each timepoint, we measured buoyant weight by placing snails on a submerged basket attached to a weight below hook and balance (Ohaus Pioneer PX224/E, Ohaus, Parsippany, NJ, USA). We then measured whole organism wet weight in air after blotting the snails dry using a paper towel. We then calculated tissue weight as the weight in air minus the buoyant weight. Weight measurements were obtained in October 2022, and in 4–5-week intervals from January – December 2023.

### 4. Feeding rate

We measured daily per-capita feeding rates at 4–6-week intervals throughout the experiment. First, snail enclosures were emptied of oysters from previous *ad libitum* feedings and fresh oysters were loaded. The week following loading (7-9 days), oyster consumption was measured by counting the number of dead oysters within each enclosure displaying drill-holes characteristic of *Urosalpinx* predation (Cheng et al. 2017). Live oysters displaying partial drill holes, or dead oysters without drill holes were excluded. The number of oysters consumed was divided by the number of days over which feeding rate was measured to provide daily per-capita oyster consumption at each timepoint. Because feeding rates were extremely low in the winter months, oyster consumption was instead measured over 14 days from December 2022 to March 2023. Once snails were placed into pairs to measure reproductive output, consumption was instead estimated per pair and halved, with the resulting per-capita estimate recorded once per pair to avoid sample size inflation. The size range of loaded oysters used to measure consumption was standardised against a reference batch of oysters, which were retained at each consumption timepoint and preserved in 96% ethanol. Later comparisons of dry tissue weight demonstrated that the average size of reference oysters (3.20 ± 0.13 mg, mean ± SE) did not vary significantly with time through the course of the experiment (ANOVA, F_1,98_ = 3.016, p = 0.086).

### 5. Reproductive output

To assess reproductive output, snail enclosures were checked for egg capsules every 1-3 days following pairing in March 2023. Our observations indicated that females typically take up to 3 days to finish laying. Therefore, enclosures containing females in the process of laying were left undisturbed until laying had finished. Subsequently, the date at which laying ceased was recorded, and clutches were removed for examination under stereomicroscopy to record both the total number of capsules laid and the number of individual embryos per capsule. Each clutch was then placed within an individual marked tea strainer and returned to its respective temperature regime to complete development. Following hatching, the number of successfully hatched juveniles was compared to the number of previously recorded embryos to determine hatching success.

### 6. Statistical analysis

We used generalized linear mixed models (GLMMs) to quantify the longitudinal effects of thermal regime (warm vs. cold) and population (NC vs. NH) on the response variables shell length (mm), specific growth rate (% g day^−1^), tissue weight (mg), and feeding rate (oysters consumed/ snail day^-1^), using the R package glmmTMB. For each response variable, we first determined the appropriate family, link function and dispersion parameters using a starting model incorporating fixed (categorical) effects of population, thermal regime, and measurement timepoint, as well as their interactions, with a random effect of snail identity to account for repeated measures. Checks were performed using diagnostic plotting of simulated residuals from maximal models using ‘DHARMa’ (Hartig 2023), posterior predictive checks using ‘performance’ (Lüdecke et al. 2021), and histogram plotting of the response variable. Justifications of model selection choices for each response variable are given in Appendix S1: Section S4. Once the appropriate GLMM structure was selected for each response, three additional random effects (Sex, reload status, and maternal line) were then added to each model and we used AIC comparison to determine whether the inclusion of any of these effects improved the final model. The random effect of snail identity was never removed from the final model based on AIC models, as it represented an integral component of our longitudinal study design. Details of which random effects were included in each final model is given in Appendix S1: Table S2.

Reproductive output was measured continually throughout the experiment, was highly heterogeneous and was characterised by complete separation of data at all but one timepoint in the cold regime. Therefore, rather than analyse the reproductive data longitudinally, we instead chose to analyse total reproductive output, quantified as the number of embryos produced by each female throughout the experiment. We also tested for differences in age and size at first reproduction in females, calculated based on the age at which the female finished laying her first clutch, and the size of the female at the nearest measurement timepoint, respectively. Lastly, we compared percentage hatching success, calculated as the proportion of embryos per capsule which successfully hatched. This data was analysed using either generalised or simple linear models, with final models chosen based on diagnostic checks as described above and as detailed in Appendix S1: Section S4. Random effects of maternal line and reload status were initially included in each model but removed based on AIC comparison, except for total reproductive output where maternal line was retained (Appendix S1: Table S2). For all models, anova tables with type-III sum of squares were produced using the ‘Anova’ function in the package ‘car’, with sidak post-hoc tests run using the package “emmeans”.

## Results

### 1. Growth metrics

There was a significant interactive effect of population, thermal regime and measurement timepoint on shell height (Wald χ2 = 113.88, p < 0.001), tissue weight (Wald χ2 = 39.69, p < 0.001), and specific growth rate (Wald χ2 = 148.59, p< 0.001; full statistical outputs in Appendix S2: Table S1). In both warm and cold thermal regimes, changes in size and weight across time were characterised by distinct periods of increase and periods of little to no change (Fig. 2B-C), closely corresponding to changes in temperature throughout the experiment (Fig. 2A). These temporal patterns were also mirrored by changes in growth rate across time, with the highest growth during the initial summer period and later during a second growth window in the second summer of the experiment (Fig. 3A).

**Figure 2:**
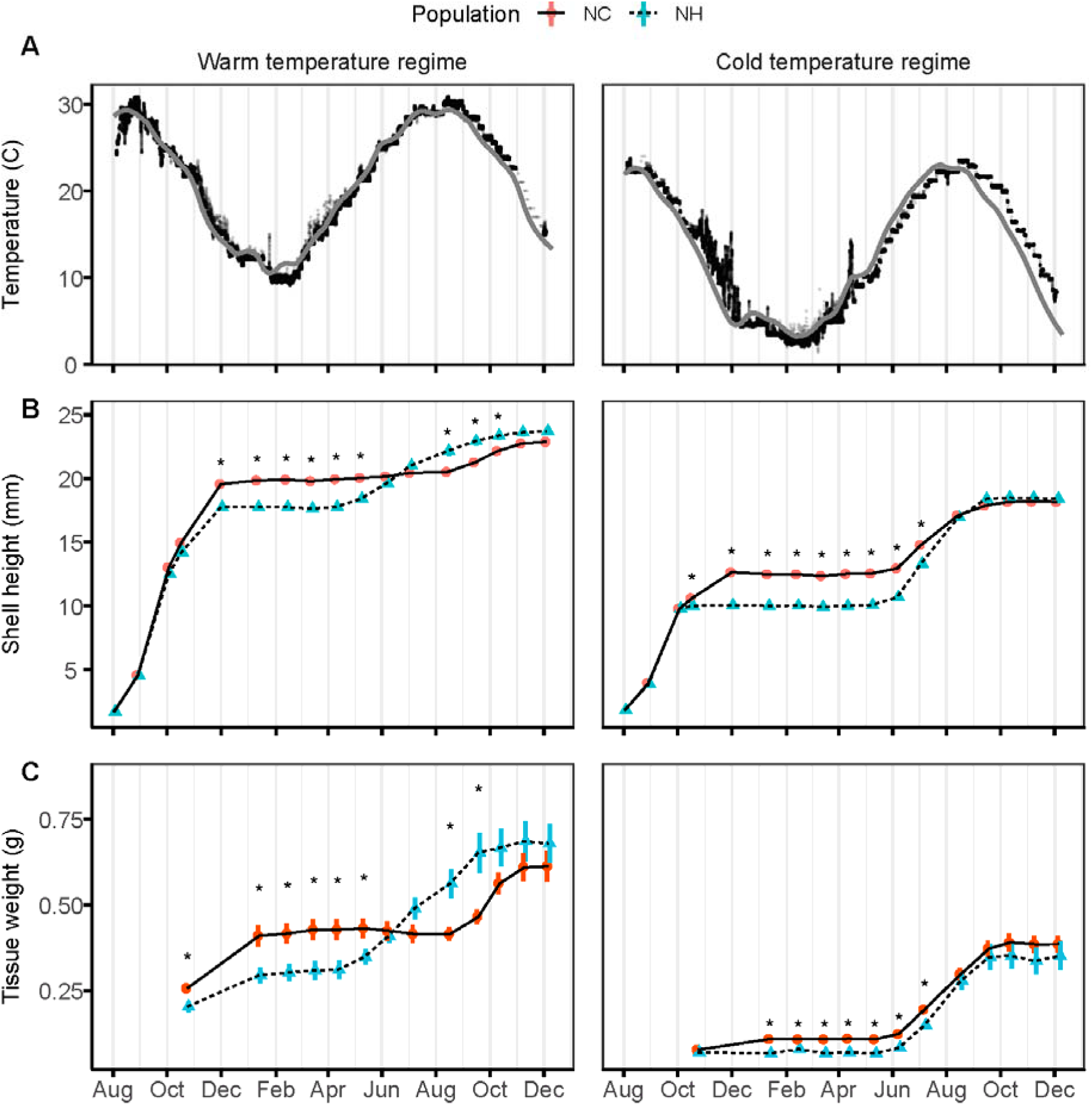
Growth trajectories in response to simulated reciprocal transplant temperature regimes in two populations of *Urosalpinx cinerea*. A) Experimental temperature profiles experienced in Warm (North Carolina) temperature regime (left panel) and Cold (New Hampshire) temperature regime (right panel). Grey lines indicate field temperature profiles used as reference to construct the two experimental temperature regimes (see Methodology 1 and Appendix S1: Section S1). B) Shell height (measured from spire to siphon tip in mm; mean ± se). C) Tissue weight (buoyant weight subtracted from whole organism wet weight, in g; mean ± se). Asterisks denote significant differences between populations within their respective temperature regime at a particular timepoint.

**Figure 3:**
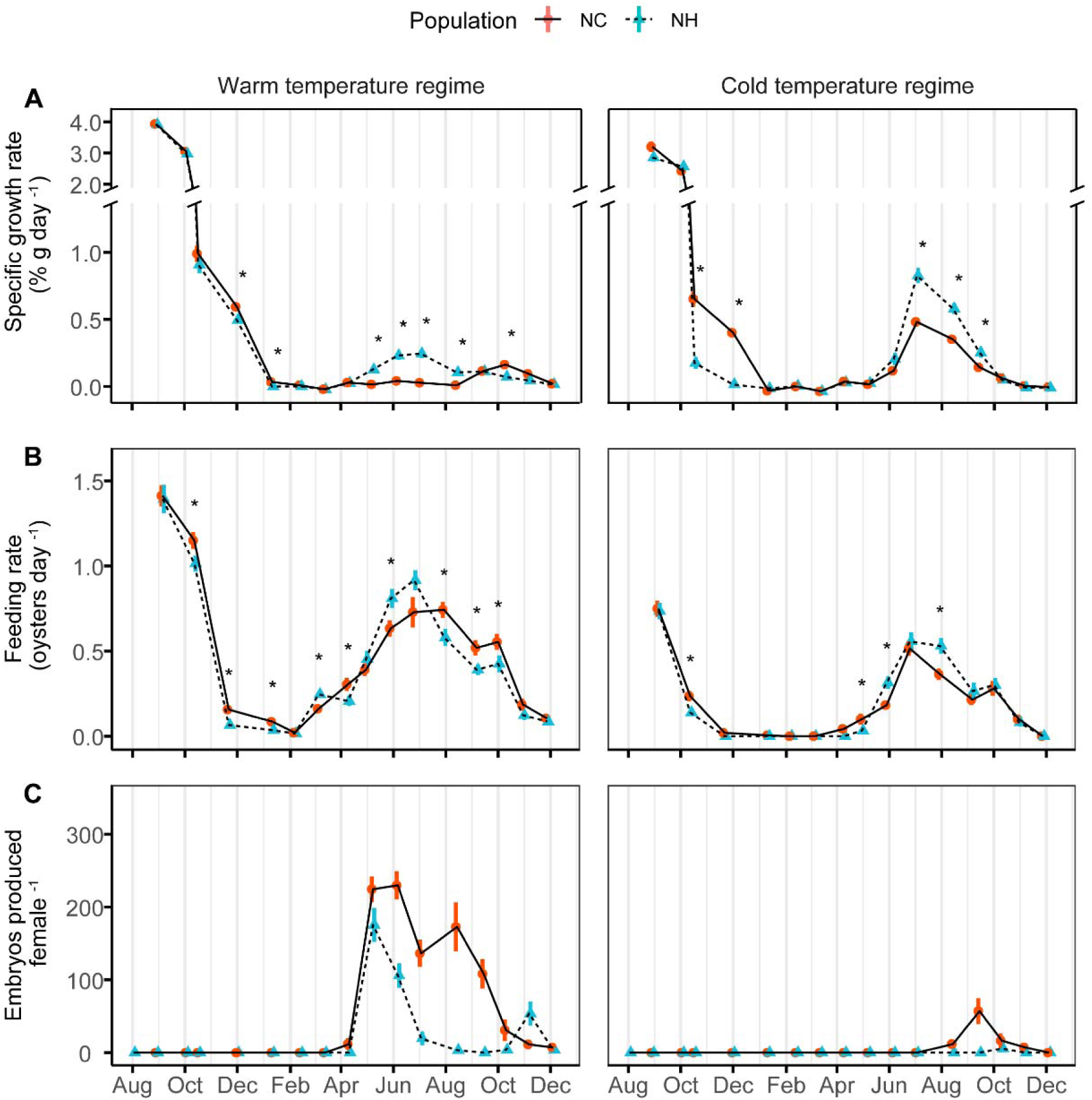
Seasonal shifts in life-history traits under simulated reciprocal transplant temperature regimes in two populations of *Urosalpinx cinerea*. Left panels: Warm (North Carolina) temperature regime. Right panels: Cold (New Hampshire) temperature regime. A) Specific growth rate (% g day-1; mean ± se; note the broken y axis). B) Feeding rate (oysters consumed / day; mean ± se). C) Reproductive output (eggs/ female; mean ± se). Asterisks denote significant differences between populations within their respective temperature regime at a particular timepoint; note that for feeding rate, post-hoc comparisons within each panel are derived from a separate statistical model as data from the two regimes was analysed separately (see methods 6.)

Across both temperature regimes, NC snails achieved a higher growth rate than NH snails in the latter part of their first summer in both temperature regimes (Fig. 3A) and subsequently possessed significantly (p<0.05) larger shell sizes and higher tissue weight during their first winter (Fig. 2B-C). However, in the warm regime a reversal of this pattern was observed during the second summer, with the NC population displaying little to no growth from May- August, while NH snails displayed higher growth in the same period evidenced by significant post-hoc comparisons between the two populations (p<0.05; Fig. 3A). This resulted in the NH snails possessing significantly larger shell heights and higher tissue weight compared to the NC population in the latter half of the second summer (Fig. 2B-C) although a late increase in growth rate in the NC population towards the end of the season in October and November (Fig. 3A) led to size differences becoming non-significant by the end of the experiment (Fig. 2B-C). In the cold regime, both populations continued to grow throughout the second summer. However, growth rates were significantly higher in the NH population from July to September (Fig. 3A), leading to no significant differences in shell height or tissue weight between the two populations in the cold regime by the end of the experiment (Fig. 2B-C).

### 2. Feeding rate

There was a significant interactive effect of population and measurement timepoint on feeding rate in both the warm regime (Wald χ2 =65.89, p < 0.001) and cold regime (Wald χ2 = 36.4, p < 0.001) (note that a longer period of zero feeding in the cold regime necessitated the separate analysis of data from the warm and cold regimes, see Appendix S1: Section S4 and Appendix S2: Table S1). In general, changes in feeding rate throughout the experiment mirrored those for growth rate (Fig. 3B; Fig. 3A). For instance, in the warm regime, feeding rates were initially higher in the NC population, corresponding to higher initial increases in size leading into the first winter, while in June and July of the second summer period feeding rates were higher in the NH population, corresponding to a period in which rapid growth was observed in this population. In the cold regime, differences between populations were less pronounced; however, a significantly higher feeding rate in the NH population was observed in August of the second summer corresponding to rapid growth in this population at this timepoint.

### 3. Reproductive traits

Reproductive output, measured as the overall total number of embryos produced per female, displayed a significant interactive effect of both population and thermal regime (Wald χ2 = 16.98, p < 0.0001). Reproductive output was higher in the warm regime and in the NC population, with an average of 932.47 ± 83.52 embryos (mean ± SE) produced in the warm regime and 92.78 ± 29.82 embryos in the cold regime, compared to 92.78 ± 29.82 and 5.69 ± 3.40 embryos respectively in the NH population (Fig. 4A). Additionally, while all females in the NC population and all but one female in the NH population reproduced in the warm regime, the number of females which did not reproduce at all in the cold regime was higher in the NH population (12/16 females) than in the NC population (10/18 females) (Fig. 4 accompanying text).

**Figure 4:**
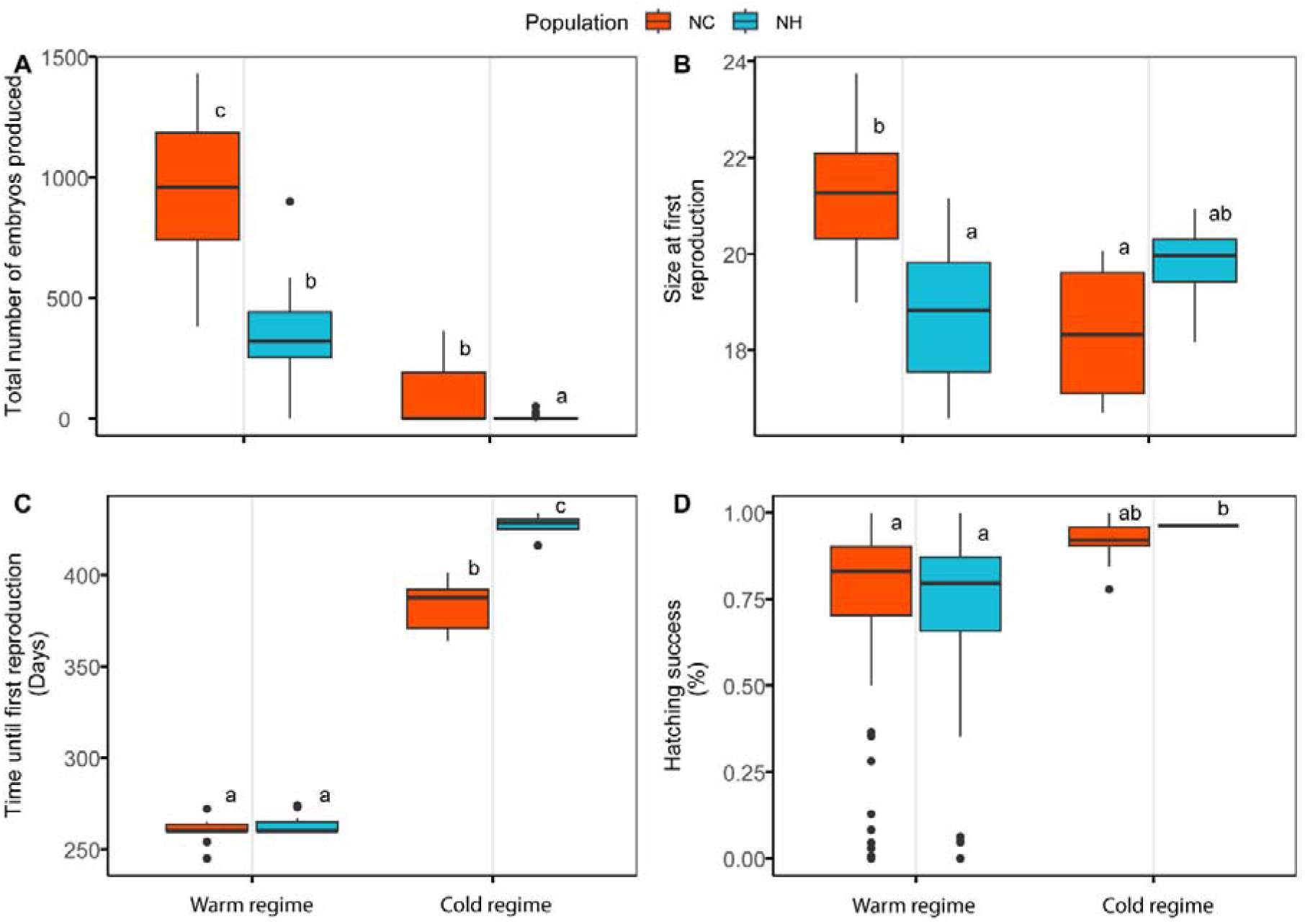
Reproductive traits under simulated reciprocal transplant temperature regimes in two populations of *Urosalpinx cinerea*. A) The total number of embryos produced per female by the end of the experiment. B) Female size at first reproduction (shell length in mm). C) Age at first reproduction in females, counted from the experiment start date. D) Percentage hatching success of offspring. Different letters indicate statistically significant post-hoc comparisons between groups (P<0.05). The number of females that had reproduced at least once vs. the total number in each treatment: population combination by the end of the experiment was as follows: warm regime, NC population: 15/15; warm regime, NH population: 13/14; cold regime, NC population: 8/18; cold regime, NH population: 4/16

Among those females which did reproduce, there were significant interactive effects of population and thermal regime on size (F_1,36_ = 12.22, p<0.001) and age (F_1,36_ = 44.371, p<0.0001) at first reproduction. In the warm regime, both populations began reproducing at near identical ages (NC - 259.53 ± 1.83 days vs. NH - 263.77 ± 1.40 days; Fig. 4C), with the NC population being significantly larger than the NH population (NC - 21.26 ± 0.37 mm vs. NH -18.82 ± 0.42 mm; Sidak post-hoc test, p<0.05; Fig. 4B). It must be highlighted that, because snails were paired in late March 2022 and onset of reproduction occurred very quickly after this in both populations in the warm regime (Fig. 2C), it is possible that snails would have been capable of reproducing at an earlier age and smaller size under natural conditions. In the cold regime, by contrast, onset of reproduction occurred well after pairing and significantly later than in the warm regime (Sidak post-hoc tests, all p<0.05), with NH snails being significantly older than NC snails (NH - 426.75 ± 3.82 days vs. NC - 383.00 ± 4.74 days; Sidak post-hoc test, p<0.05; Fig. 4C). NH snails also trended towards larger size at first reproduction (NH - 19.76 ± 0.58 mm vs. NC - 18.35 ± 0.50 mm), although this difference was not significant (Sidak post-hoc test, p>0.05; Fig. 4B). There was no significant interactive effect of population and temperature regime on percentage hatching success (Wald χ2 = 0.29, p = 0.59). When the interaction term was excluded from the model, hatching success differed significantly across temperature regimes (Wald χ2 = 13.11, p < 0.001) but not between populations (Wald χ2 = 1.18, p=0.28), being lower in the warm than cold regime (Fig. 4D).

## Discussion

### Responses are consistent with Pace-of-Life divergence across thermal environments

Understanding the mechanisms driving local adaptation of life-history strategies can provide crucial insights into forecasting species responses to warming but remains poorly studied within the context of environmentally relevant temperature regimes and exposure timescales. Our reciprocal transplant experiment, which closely simulated seasonal temperature variation experienced in the field over a multiyear period, revealed a radical reshaping of life history strategies across latitudinally-distinct populations. The low-latitude NC population initially achieved a larger size than the high-latitude NH population, but this pattern switched in the second year, with the NC population instead displaying lower growth than the NH population but higher reproductive output, suggesting an early energetic switch to reproductive investment at the expense of continued growth. Although this pattern was most apparent in the warm thermal regime, perhaps due to elevated temperatures accelerating life history progression, differences in responses between the two populations were consistent across both temperature regimes. This contrasts with the Local Advantage scenario (Fig. 1A-C) which predicts growth in each regime should be highest in the respective locally adapted population. Instead, our results appear consistent with the Pace of Life Syndrome hypothesis (POLS; Fig. 1G-I), in which local adaptation of life-history strategies across thermal environments follows a fast-slow axis. In this context, we interpret the warm-adapted NC population as having been selected for a fast strategy of high initial growth and reproductive output, with the cold adapted NH population instead pursuing a slow strategy of slower but sustained growth and delayed reproduction. Our study highlights the potential limitations of short-term comparisons of growth without factoring in seasonal temperature variation, as responses varied with time during our experiment, demonstrating a multifaceted interaction between temperature, growth and life history strategies in determining population-level responses across latitude.

A fast pace of life at low latitudes may represent an evolved response to environmental constraints imposed by warm environments, including increased predation risk (Freestone et al., 2021; Walker et al., 2020) and elevated thermal mortality risk (Verberk et al., 2021). As these constraints limit life expectancy, rapid growth to achieve early, high reproductive output may be favoured. At the same time, potential deleterious effects of rapid growth on performance and longevity in later life, such as increased oxidative damage, may be less relevant, leading to a “grow fast, die young” strategy (Metcalfe & Monaghan, 2003; Réale et al., 2010). Indeed, field trials conducted in parallel to the present study measuring relative predation rates on *Urosalpinx* have revealed over 14-fold greater predation risk in North Carolina as in New Hampshire, confirming that predation pressure may represent a crucial selective agent driving high reproductive investment in the southern population (Rawson et al., *manuscript in preparation*). In addition, metabolic adaptation in ectotherms is potentially constrained by thermodynamic effects of temperature variation across latitude, potentially leading to selection for faster growth strategies in warm environments due to increased biological rates (Angilletta et al., 2010; Frazier et al., 2006). Conversely, fecundity scales more greatly with size in cold environments (Weetman and Atkinson 2004, Arendt 2015), potentially favouring genotypes which delay reproduction until a larger size is achieved. In addition to physiological and life-history traits, Pace-of-Life divergence is commonly associated with personality traits, with fast-paced individuals possessing higher boldness and risk-taking behaviour (Dammhahn et al., 2018; Réale et al., 2010). While we did not measure behavioural responses in this study, data from the aforementioned parallel study indicates strong divergence in behavioural traits between the two populations in accordance with the fast-slow axis, with the low latitude NC snails displaying higher movement speed and increased boldness compared to snails originating from the NH population (Rawson et al., *manuscript in preparation*).

Factors affecting the directionality of Pace-of-life responses across latitude

Previous studies that have found support for a faster pace-of-life in low latitude populations have focused on short-lived species, including damselfly larvae, which exhibit increased risk-taking, faster growth, and higher metabolic rates (Debecker and Stoks 2019, Tüzün and Stoks 2022), and ants, which show heightened exploratory and foraging behaviours (Segev et al. 2017). By contrast, a novel aspect of our study is that it provides evidence for POLS divergence of life-history traits in a relatively long-lived invertebrate ectotherm, *Urosalpinx*, which may potentially live up to 10–15 years in the wild (Carriker 1955). Comparison with patterns seen in other species suggests that differences in the directionality of POLS responses across latitude may be linked to the differing extents to which growth windows are temporally constrained across taxa. For instance, although our findings apparently conflict with the CnGV model (Fig 1D-F), which posits that cold-adapted populations are selected for rapid growth to mitigate against reduced season length (Conover et al. 2009), this disparity may hinge on the key assumption of CnGV that life-history constraints force high-latitude populations to achieve comparable growth to low-latitude populations within a similar timeframe. For instance, in a classic example of CnGV, the Atlantic silverside *Menidia menidia*, individuals have a lifespan of under two years and are under strong selective pressure to reach adult size by their first winter to reproduce immediately afterwards (Yamahira and Conover 2002). Thus, high latitude populations display faster growth and higher metabolic rate than low latitude populations, coupled with higher food consumption and growth efficiency (Conover and Present 1990, Present and Conover 1992). However, season length constraints at high latitude may not favour rapid maturation in long-lived species such as *Urosalpinx*, which reproduces multiple times in its life and can overwinter at immature size (as observed in our cold regime). Interestingly, *Urosalpinx* populations living at different latitudes may differ in their generation time, similar to other species, including damselflies, which also display Pace-of-life divergence across latitude (Debecker and Stoks 2019, Tüzün and Stoks 2022). Based on our results, it appears likely that the NC population could achieve maturity and reproduce within a single growing season in the field, especially given that snails in the warm regime had already achieved the body size at which they were paired in March 2023 by the end of the previous year (Figure 2B). Indeed, maturation of offspring could occur even earlier in the field given that the first eggs are produced in the spring (Carriker 1955). Conversely, the delayed onset of reproduction we observed in the cold regime (especially apparent in the NH population) may suggest that northern populations require at least two growing seasons to reach maturity.

We note that inferring latitudinal adaptation from a two-population comparison carries caveats, as observed differences between any two populations could also be due to site-specific differences (Garland and Adolph 1994). While we focused upon only two populations (NC and NH), we also collected data from an additional low-latitude population (GA) which, while limited by reduced sample size and lack of replication of maternal lines, presents an informative comparison with our main dataset (Appendix 2: Supplement S1 and Figures S1-S2). In general, growth and feeding rate in GA snails closely resembled NC snails, suggesting that the GA population is similarly selected for fast growth in accordance with faster biological rates at low latitudes. By contrast, total reproductive output in GA was intermediate between the NC and NH populations in the warm regime. This is supported another set of parallel reproductive output experiments suggesting that under local temperature conditions, the GA population has a reduced reproductive season, with a hiatus during peak summer temperatures that appear to reduce hatching success of developing embryos (Rugila et al., *in review, preprint available*). Such differences may reflect regional variation in selective pressures at different sites, such as predator abundance, which may alter specific pace-of-life responses in individual populations. Alternatively, elevated thermal sensitivity in juveniles compared to adults (Zippay and Hofmann 2010, Truebano et al. 2018) may impose a ceiling on the number of embryos produced in populations experiencing the warmest temperatures, forcing greater investment per embryo or a suspension of reproduction in the hottest part of the year. Further work is necessary to understand how traits co-evolve in response to environmental stress gradients and across different taxa, and the extent to which constraints imposed by certain environments (particularly at range edges) may alter or disrupt these correlated responses (Montiglio et al. 2018).

Common garden vs. reciprocal transplant designs

Our use of fluctuating temperature regimes, designed to reflect seasonal variation, contrasts with many studies examining latitudinal divergence in life-history traits which use comparisons under constant temperatures and common-garden approaches (Kawecki and Ebert 2004, Conover et al. 2009, De Frenne et al. 2013). While such approaches are valuable for revealing underlying genetic adaptation that may be masked under field conditions, an inherent consequence is that they risk divorcing observed responses from the environmental context under which they evolved (Kawecki and Ebert 2004). Such limitations may be particularly pronounced in studies focusing on divergence in thermal optima, as optimal temperatures often do not reflect those at which most growth and reproduction (and therefore selection) take place (Deutsch et al. 2008, Martin and Huey 2008, Dwane et al. 2023). This may explain why our findings differ from a previous study, which identified CnGV in growth rate of *Urosalpinx* with north populations displaying higher growth at the thermal optimum than southern populations (Villeneuve et al. 2021a). Crucially, the prior study used a common-garden design focusing on maximum performance and thermal optima as target traits, the latter occurring at around 27-30°C, and was restricted to juvenile snails maintained for 21 days. By contrast, our fluctuating regimes only reached such extremes for a few months each year in the warm regime. As average temperatures were thus cooler, divergence in growth between the two populations likely reflected performance at the lower end of the thermal performance curve, while our 17-month timeframe enabled longer-term growth patterns to emerge, potentially explaining why interpopulation differences observed in our study differed from those seen in the previous study. Interestingly, at early timepoints in our study - when temperatures were close to the thermal optimum - growth and body size were indeed similar across populations (Fig. 2, Fig. 3A), with NC only overtaking NH in size as temperatures declined in the boreal autumn. NH snails also exhibited higher growth during the second summer in both regimes, when temperatures also peaked (Fig. 2A). This could reflect a greater ability of the NH population to capitalise on short windows of favourable growing conditions in their home environment, consistent with the conclusions of the previous study that CnGV in maximum growth rate is an adaptation to seasonality across latitude (Yamahira and Conover 2002, Villeneuve et al. 2021a). Nonetheless, our broader finding of a faster-paced life in low latitude populations demonstrates the limitations of extrapolating latitudinal patterns from short-term common-garden designs, highlighting the importance of understanding local adaptation in the context of dynamically fluctuating temperatures and appropriate timescales (Lovell et al. 2023).

Shifts towards a faster pace of life under climate change

The divergence in life-history strategies observed in our study has important implications for understanding and forecasting future response to climate change across species’ ranges. In our experiment, the high-latitude NH population achieved accelerated growth and reproductive output in the warm regime compared to the cold regime, illustrating that a shift to warmer average temperatures under climate change may lead to plastic and elevated performance in certain traits (but not necessarily fitness) even in the absence of future adaptation (Deutsch et al. 2008). This corresponds with previous studies indicating that *Urosalpinx* populations currently living in colder habitats will experience shifts towards their thermal optimum for growth under warming (Cheng et al. 2017, Villeneuve et al. 2021a). Moreover, our finding that existing local adaptation to warm environments is associated with a faster pace-of-life suggests that future shifts in phenotypic responses could act synergistically with warming, leading to the effects of an accelerated life history under climate change being further exacerbated by directional selection. This contrast sharply with expectations under CnGV, previously proposed as a major mechanism by which ectotherms may respond to climate change (Conover et al. 2009, Baumann and Conover 2011). Under CnGV, adaptive responses would act to mask environmental effects on the phenotype, leading to no overall shift in a population’s performance under increased temperatures (Skelly 2010, Stoks et al. 2014). By contrast, if selection under warming favours an accelerated pace of life, this could compound impacts on stressed communities already undergoing restructuring under climate change (Vergés et al. 2019), with potentially drastic consequences for ecosystem function. As Pace-of-life responses are also driven by factors such as predation pressure, the assumption that climate change responses of species will mirror existing latitudinal gradients is contingent on the assumption that their respective predators will shift polewards, or increase in abundance, as temperatures increase. For *Urosalpinx*, this is plausible, as a major predator of *Urosalpinx* at low latitudes, the blue crab *Callinectes sapidus*, is exhibiting northwards range shifts (Johnson 2015). More broadly, temperate ecosystems globally are undergoing range shifts of mobile species towards higher latitudes, leading to fundamental shifts in the character of communities to resemble warmer locations via “tropicalization” (Vergés et al. 2019). For less dispersive species, including *Urosalpinx*, it is likely that selective environments today experienced by high latitude populations will come to increasingly resemble those today experienced by lower latitude populations, with concomitant changes in selective pressures. This suggests that these high latitude populations would be selected for faster initial growth rate, early maturation, and greater fecundity. In contrast, low latitude populations that experience warming may be selected for greater growth rate up until maximum trait performance begins to decline, possibly resulting reduced hatchling survival and reproductive output, ultimately resulting in local extinction (Villeneuve, Komoroske, and Cheng 2021a; Rugila et al., *in review, preprint available*).

## Conclusions

The impacts of warming on marine ecosystems will be considerable, likely leading to a shifting seascape of life history strategies. Determining whether latitudinal responses of individual species generally follow responses consistent with the POLS, or countergradient responses, is critical to understanding whether local adaptation will act synergistically with, or counteract, the effects of climate change at the community level. If the directionality of life-history responses across latitude is highly dependent on intraspecific differences in life-history strategies, longevity, and predator-prey interactions, this indicates urgent work is needed to characterise the nature of these responses across a wide array of taxa, to enhance future predictions of how functional dynamics of ecosystems will be affected under warming. Our results imply that high-latitude populations of marine predators could respond to, and in so doing compound, ecosystem-wide community shifts through a radical reshaping of life-history strategies, potentially adopting a faster-paced strategy currently seen in low-latitude populations. These findings underscore the importance of considering local adaptation of life-history strategies and the need to understand responses within the context of environmental temperature fluctuations and biologically relevant time scales experienced in nature.

## Supporting information

Appendix S2 results supplement

## Acknowledgements

The authors would like to thank Jeb Byers and Joshua Lord for assistance with identifying collection sites, and Estefany Argueta for assistance with animal husbandry

