## Appendix S1 methods supplement for "Adaptation to warm environments with a fast pace of life in a marine predatory snail"

### Appendix S1: Methodology supplement

#### Section S1: Maintenance of temperature regimes

To inform the environmental simulation experiment, we obtained reference temperature data for the warm temperature regime from NOAA National Data Buoy Center (NDBC) station BFTN7, located 0.1 km from the sample site where the North Carolina (NC) *Urosalpinx* population was sourced (34°43'05"N 76°40'16"W). For the cold regime, we sourced reference temperature data from the National Estuarine Research Reserve System (NERRS) station GRBGBWQ, located 1.97 km from the New Hampshire (NH) sample site within Great Bay (43°05'20" N 70°51'58" W). However, as data collection from this station ceases between the 14^th^ December and the 5^th^ April each year due to ice formation, we substituted reference temperature data for this winter period with data from NDBC station CMLN3, owned by the University of New Hampshire. This station is located at the mouth of Great Bay (43°4'12" N 70°42'0" W) 13.6 km from the sampling location. We believe this is a reasonable substitution because winter temperatures are spatially homogenous within the estuary. Reference temperatures were obtained by averaging daily mean temperatures for the years 2019 – 2021.

Typically, temperatures in the two temperature regimes were maintained ± 1°C of setpoint temperatures; however, deviations in temperatures did occur during the experiment primarily due to limitations in the ability to maintain temperatures in the building containing the seawater tables during heatwaves (Fig. 2A). However, local deviations from seasonal temperature averages are known to occur in the field in intertidal environments (Helmuth et al. 2006). Temperature spikes during the summer were mitigated against by the addition of frozen seawater bottles to the sumps.

In addition, because winter temperature in the cold regime required temperatures of below 5°C to be maintained which could not be safely maintained by the chiller units, from the 23^rd^ January until the 20^th^ March 2023 the snails from both regimes were instead kept inside 150 L plastic storage containers inside separate 200 L chest freezers (Magic Chef 7.0, Magic Chef, Wood Dale, IL, USA), which were connected to thermostat units (NEMA 4X, AquaLogic Inc., Monroe, NC, USA) to maintain setpoint temperatures for the respective regimes. These contained seawater and biological and mechanical filtration and were checked daily to maintain water quality to the same parameters as described in the main text and below.

#### Section S2: Animal care

Water quality during the reciprocal transplant experiment was measured daily using a handheld multiparameter probe (YSI Professional Plus, YSI Incorporated, Yellow Springs, OH, USA) to maintain salinity (30 ± 1 PSU) and temperature (± 1°C of setpoint). We also measured ammonia twice per week using ammonia test kits (Red Sea Fish Pharm Ltd., Herzliya, Israel). Oysters that were fed to snails were maintained inside aerated plastic bins within flow-through seawater systems. Oysters were fed five times a week with Shellfish Diet at a target concentration of 0.4 mg dry weight algae oyster^-1^ (Reed Mariculture, Campbell, CA, USA).

#### Section S3: Snail pairing

Pairing of most individuals occurred on the 20^th^ of March 2023. For the NC and NH populations, females were outcrossed with experimental males from a different maternal line, while for the GA population (which were from a single maternal line) females were instead paired with available sibling males. Because there was an excess of females in the NC population in the cold regime, and in the GA population in both the warm and cold regimes, these females were paired with wild males which had been collected from the same field locations as the original broodstock.

As of the 20^th^ of March, a total of 11 individuals in the NH population in the cold regime were below the size of the smallest snail positively identified as a male at the time of pairing (10 mm). These could therefore not be reliably sexed, so were instead paired with other putatively immature snails from the same population. On the 10^th^ of July 2023, these snails had grown sufficiently to be sexed, and were either paired with each other, unpaired experimental males from the NH population transferred from the warm regime (which were subsequently excluded from further data collection beyond this date, n=6), or additional wild males (n=3).

Because the feeding rate of field-collected or transplanted males could not be distinguished from that of experimental females with which they were paired, consumption data from any pairs containing these males was excluded from data analysis, although size data and fecundity data from the paired females was still used.

Odd individuals which could not be paired, or individuals whose partner died during the experiment, were kept inside the same sized enclosures but fed half the allocation of oysters fed to paired snails within their temperature regime. These individuals were retained to provide continuous growth and consumption data.

#### Section S4: GLMM model selection

For shell length, data was largely normally distributed, but variance increased with the mean, so a Gaussian error distribution with log link was used. For specific growth rate, data was strongly right skewed and contained many values at or close to zero, representing winter periods where little growth occurred. In addition, negative calculated values for growth rate were present in the dataset. These were converted to zero prior to analysis, both to simplify model selection, and in recognition of their likely status as artifacts of measurement error during period of no growth. Following this, a Tweedie distribution, suitable for highly zero-inflated data , was used, combined with a dispersion formula allowing variances to differ both by regime and by measurement timepoint. For tissue weight, data was non-normal, positive and continuous, so a Gamma distribution with log link function was used.

For feeding rate, data was positive, continuous, and characterised by high variation between measurement timepoints. Complicating matters, at five timepoints within the cold regime only (2,3,4, and 13) consumption rates of zero were observed across all individuals across one or both populations, resulting in complete separation at these timepoints and precluding the use of zero-inflated or hurdle-type approaches. To account for this, separate GLMM models were run for the warm and cold regimes, with the five aforementioned timepoints excluded from the model for the cold regime but included for the warm regime (where complete separation did not occur). Both models used gaussian distributions and identity link functions with a dispersion formula allowing variances to differ across measurement timepoints.

Reproductive output data per female, measured cumulatively over the entire experiment, was right-skewed and zero-inflated, accounting for multiple females which had zero reproductive output throughout the experiment. We therefore used a zero-inflated generalized linear model with negative binomial family and log link function, with temperature regime and population as categorical predictor variables. Random effects of maternal line and reload status were initially included but removed based on AIC comparison. For both age and size at first reproduction in females, a gaussian distribution with identity link function was found to be most suitable, with all random effects being excluded during model selection; thus, final models were run as linear models in base R. Hatching success, quantified as the proportion of embryos per capsule which successfully hatched, ranged in value freely between zero and one; therefore, a generalized linear model with quasibinomial family and identity link was run using the “glm” function found in the “stats” package.

Table S1.: Snail mortality throughout the experiment, excluding individuals which died during the first three weeks and were replaced; see Methodology 2.

| Regime | Alive at end of experiment | Population | | |
| --- | --- | --- | --- | --- |
|  |  | NC | NH | GA |
| Warm | *Yes* | 31 | 36* | 16 |
|  | *No* | 9 | 4 | 4 |
| Cold | *Yes* | 28 | 23 | 9 |
|  | *No* | 12 | 17 | 11 |

*This includes six male individuals which were transferred to the cold regime on the 10^th^ July 2023, in order to provide mates for NH females as noted below. Data from these individuals was included in the dataset up until the point of transfer.

Table S2: Random effects model selection table

The random effect structure used in final analysis for each variable is given in bold. “NA” values indicate models which failed to converge. Random effects are as follows: Snail ID = individual snail identity, Sex = male or female, Maternal Line – identity of the parent female for each snail, Reload status = whether the snail was added as a replacement to compensate for mortality within the first three weeks; see methods 2.

| *Random effect structure* | *Df* | *AIC* | *logLik* | *∆AIC* |
| --- | --- | --- | --- | --- |
| **Shell height** |  |  |  |  |
| **(1\|Snail ID)+ (1\|Sex)+ (1\|Maternal Line)** | **72** | **5713.99** | **-2785.00** | **0** |
| (1\|Snail ID)+ (1\|Sex)+ (1\|Reload status)+ (1\|Maternal Line) | 73 | 5715.99 | -2785.00 | 2 |
| (1\|Snail ID)+ (1\|Sex) | 71 | 5716.02 | -2787.01 | 2.03 |
| (1\|Snail ID)+ (1\|Sex)+ (1\|Reload status) | 72 | 5718.02 | -2787.01 | 4.03 |
| (1\|Snail ID) | 70 | 5721.58 | -2790.79 | 7.58 |
| (1\|Snail ID)+ (1\|Reload status) | 71 | 5723.58 | -2790.79 | 9.58 |
| (1\|Snail ID)+ (1\|Maternal Line) | 71 | NA | NA | NA |
| (1\|Snail ID)+ (1\|Reload status)+ (1\|Maternal Line) | 72 | NA | NA | NA |
| **Growth rate** |  |  |  |  |
| **(1\|Snail ID)+ (1\|Sex)+ (1\|Reload status)** | **100** | **-840.48** | **520.24** | **0** |
| (1\|Snail ID)+ (1\|Sex)+ (1\|Maternal Line) + (1\|Reload status) | 101 | -838.48 | 520.24 | 2 |
| (1\|Snail ID)+ (1\|Reload status) | 99 | -837.90 | 517.95 | 2.58 |
| (1\|Snail ID)+ (1\|Reload status)+ (1\|Maternal Line) | 100 | -835.90 | 517.95 | 4.58 |
| (1\|Snail ID)+ (1\|Sex) | 99 | -832.44 | 515.22 | 8.03 |
| (1\|Snail ID)+ (1\|Sex)+ (1\|Maternal Line) | 100 | -830.44 | 515.22 | 10.03 |
| (1\|Snail ID) | 98 | -828.29 | 512.14 | 12.19 |
| (1\|Snail ID)+ (1\|Maternal Line) | 99 | -826.29 | 512.14 | 14.19 |
| **Tissue weight** |  |  |  |  |
| **(1\|Snail ID)+(1\|Sex)** | **55** | **-1458.47** | **784.24** | **0** |
| (1\|Snail ID)+(1\|Sex) + (1\|Maternal Line) | 56 | -1457.46 | 784.73 | 1.01 |
| (1\|Snail ID)+(1\|Sex) +(1\|Reload status) | 56 | -1456.47 | 784.24 | 2 |
| (1\|Snail ID) | 54 | -1455.47 | 781.73 | 3.01 |
| (1\|Snail ID)+(1\|Sex) + (1\|Maternal Line)+(1\|Reload status) | 57 | -1455.46 | 784.73 | 3.01 |
| (1\|Snail ID)+(1\|Maternal Line) | 55 | -1455 | 782.5 | 3.47 |
| (1\|Snail ID)+(1\|Reload status) | 55 | -1453.47 | 781.73 | 5.01 |
| (1\|Snail ID)+(1\|Maternal Line)+(1\|Reload status) | 56 | -1453 | 782.5 | 5.47 |
| **Consumption (Warm regime)** | | | |  |
| **(1\|Snail ID)** | **46** | **-695.72** | **393.86** | **0** |
| (1\|Snail ID)+(1\|Reload status) | 47 | -694.4 | 394.2 | 1.32 |
| (1\|Snail ID)+(1\|Maternal Line) | 47 | -693.72 | 393.86 | 2 |
| (1\|Snail ID)+(1\|Reload status)+(1\|Maternal Line) | 48 | -692.45 | 394.22 | 3.28 |
| **Consumption (Cold regime)** |  |  |  |  |
| **(1\|Snail ID)** | **28** | **-237.47** | **146.73** | **0** |
| (1\|Snail ID)+(1\|Reload status) | 29 | -235.47 | 146.73 | 2 |
| (1\|Snail ID)+(1\|Maternal Line) | 29 | -235.47 | 146.73 | 2 |
| (1\|Snail ID)+(1\|Reload status)+(1\|Maternal Line) | 30 | -233.47 | 146.73 | 4 |
| **Size at first reproduction** |  |  |  |  |
| **none** | **5** | **148.44** | **-69.22** | **0** |
| (1\|Maternal Line) | 6 | 149.75 | -68.88 | 1.31 |
| (1\|Reload status) | 6 | 150.44 | -69.22 | 2 |
| (1\|Maternal Line)+(1\|Reload status) | 7 | 151.75 | -68.88 | 3.31 |
| **Age at first reproduction** |  |  |  |  |
| **none** | **5** | **288** | **-139** | **0** |
| (1\|Reload status) | 6 | 290 | -139 | 2 |
| (1\|Maternal Line) | 6 | 290 | -139 | 2 |
| (1\|Maternal Line)+(1\|Reload status) | 7 | 292 | -139 | 4 |
| **Total reproductive output** |  |  |  |  |
| **(1\|Maternal Line)** | **7** | **600.01** | **-293** | **0** |
| (1\|Reload status) | 7 | 601.28 | -293.64 | 1.27 |
| (1\|Egg_tag)+(1\|Is_Replacement) | 8 | 602.01 | -293 | 2 |
| none | 6 | 610.19 | -299.1 | 10.19 |
