## Appendix S2 results supplement for "Adaptation to warm environments with a fast pace of life in a marine predatory snail"

### Table S1: Type 3 ANOVA tables for all statistical tests conducted in the study.

| **Shell height** | | | |  |
| --- | --- | --- | --- | --- |
| *term* | *Χ^2^* | *df* | *P* |  |
| (Intercept) | 10712.83 | 1 | <0.001 |  |
| Regime | 600.49 | 1 | <0.001 |  |
| Population | 10.03 | 1 | <0.001 |  |
| Measure | 26307 | 16 | <0.001 |  |
| Regime:Population | 7.71 | 1 | 0.01 |  |
| Regime:Measure | 2992.7 | 16 | <0.001 |  |
| Population:Measure | 1101.31 | 16 | <0.001 |  |
| Regime:Population:Measure | 113.88 | 16 | <0.001 |  |
| *Random effect* |  |  | *Variance* |  |
| Snail ID |  |  | 0.0042627 |  |
| Sex |  |  | 0.001019 |  |
| Maternal Line |  |  | 0.0005703 |  |
| **Growth rate** | | | |  |
| *term* | *Χ^2^* | *df* | *P* |  |
| (Intercept) | 1196.32 | 1 | <0.001 |  |
| Regime | 4.14 | 1 | 0.04 |  |
| Population | 3.09 | 1 | 0.08 |  |
| Measure | 13017.28 | 15 | <0.001 |  |
| Regime:Population | 0.3 | 1 | 0.58 |  |
| Regime:Measure | 614.83 | 15 | <0.001 |  |
| Population:Measure | 257.91 | 15 | <0.001 |  |
| Regime:Population:Measure | 148.59 | 15 | <0.001 |  |
| *Random effect* |  |  | *Variance* |  |
| Snail ID |  |  | 2.173E-11 |  |
| Sex |  |  | 0.000964 |  |
| Reload status |  |  | 0.005747 |  |
| **Tissue weight** | | | |  |
| *term* | *Χ^2^* | *df* | *P* |  |
| (Intercept) | 305.76 | 1 | <0.001 |  |
| Regime | 363.38 | 1 | <0.001 |  |
| Population | 11.12 | 1 | <0.001 |  |
| Month | 5155.08 | 12 | <0.001 |  |
| Regime:Population | 3.41 | 1 | 0.06 |  |
| Regime:Month | 1020.25 | 12 | <0.001 |  |
| Population:Month | 209.09 | 12 | <0.001 |  |
| Regime:Population:Month | 39.69 | 12 | <0.001 |  |
| *Random effect* |  |  | *Variance* |  |
| Snail ID |  |  | 0.02447 |  |
| Sex |  |  | 0.01094 |  |
| **Consumption (Warm regime)** | | | |  |
| *term* | *Χ^2^* | *df* | *P* |  |
| (Intercept) | 3455.35 | 1 | <0.001 |  |
| Population | 3.08 | 1 | 0.86 |  |
| Measure | 3168.49 | 14 | <0.001 |  |
| Population:Measure | 65.89 | 14 | <0.001 |  |
| *Random effect* |  |  | *Variance* |  |
| Snail ID |  |  | 0.0002321 |  |
| **Consumption (Cold regime)** | | | |  |
| *term* | *Χ^2^* | *df* | *P* |  |
| (Intercept) | 277.8 | 1 | <0.001 |  |
| Population | 0.04 | 1 | 0.85 |  |
| Measure | 274.25 | 8 | <0.001 |  |
| Population:Measure | 36.4 | 8 | <0.001 |  |
| *Random effect* |  |  | *Variance* |  |
| Snail ID |  |  | 0.03589 |  |
| **Total reproductive output** | | | |  |
| *term* | *Χ^2^* | *df* | *P* |  |
| (Intercept) | 898.39 | 1 | <0.001 |  |
| Regime | 74.09 | 1 | <0.001 |  |
| Population | 43.53 | 1 | <0.001 |  |
| Regime:Population | 16.98 | 1 | <0.001 |  |
| **Hatching success** | | | |  |
| *term* | *Χ^2^* | *df* | *P* |  |
| Regime | 3.01 | 1 | 0.08 |  |
| Population | 0.00 | 1 | 0.24 |  |
| Regime:Population | 0.29 | 1 | 0.59 |  |
| **Size at first reproduction** | | | | |
| *term* | *SS* | *df* | *F* | *P* |
| (Intercept) | 11863.21 | 1 | 5725.27 | <0.001 |
| Regime | 6.29 | 1 | 3.03 | 0.09 |
| Population | 2.86 | 1 | 1.38 | 0.25 |
| Regime:Population | 25.37 | 1 | 12.24 | <0.001 |
| Residuals | 74.05 | 36 |  |  |
| **Age at first reproduction** | | | | |
| *term* | *SS* | *df* | *F* | *P* |
| (Intercept) | 3426657 | 1 | 50499.47 | <0.001 |
| Regime | 158221.7 | 1 | 2331.75 | <0.001 |
| Population | 4440.21 | 1 | 65.44 | <0.001 |
| Regime:Population | 3010.79 | 1 | 44.37 | <0.001 |
| Residuals | 2442.79 | 36 | NA | NA |

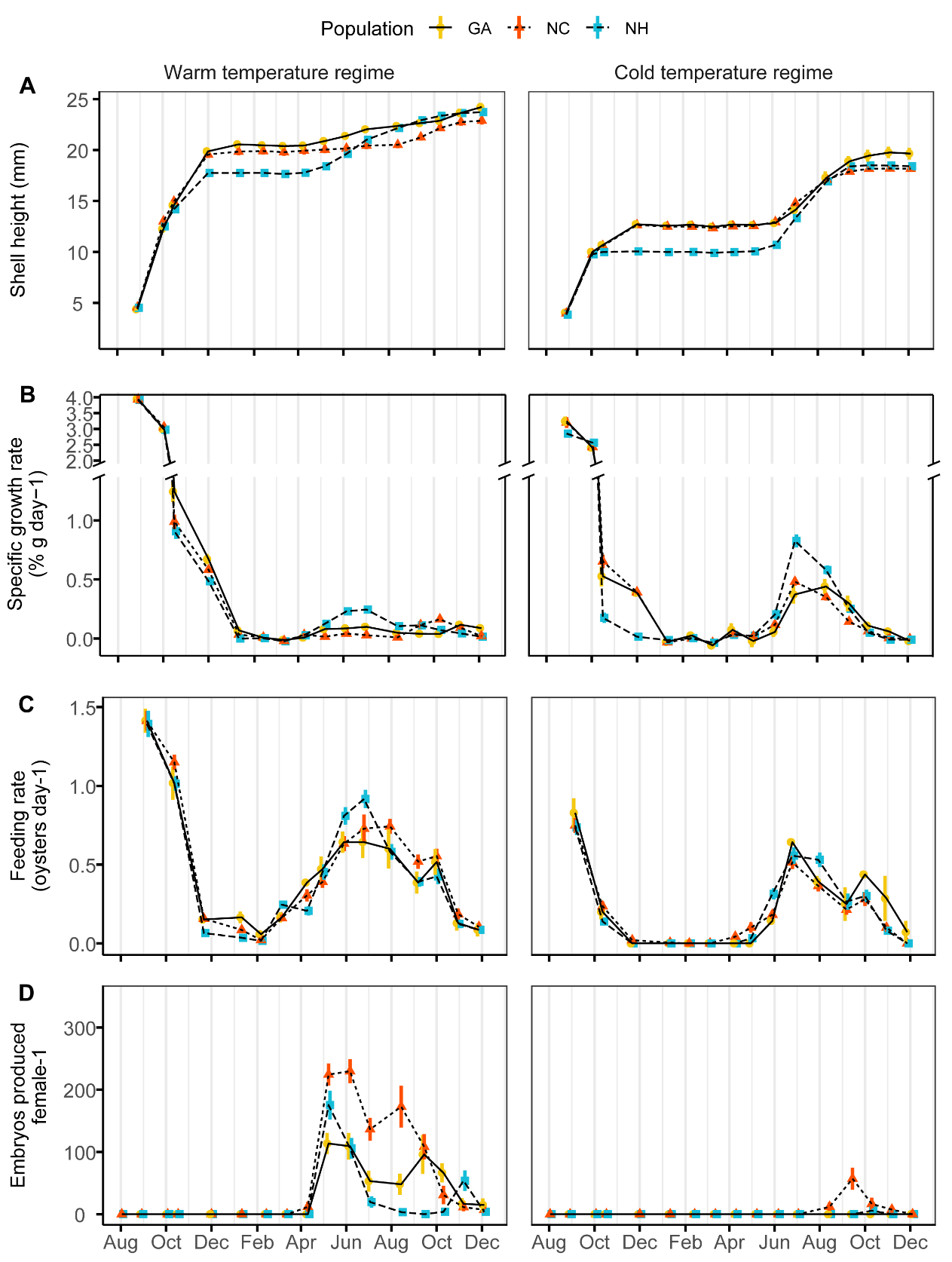

### Figure S1:

### Longitudinal experimental data incorporating the Georgia population. A) Shell height, B) Growth, C) feeding rate and D) reproductive output in all three populations including Georgia (gold), which was omitted from the main study (see Methods 1). Graphics for each panel depict measurements as described in the captions for Figure 2B, 3A, 3B and 3C, respectively.

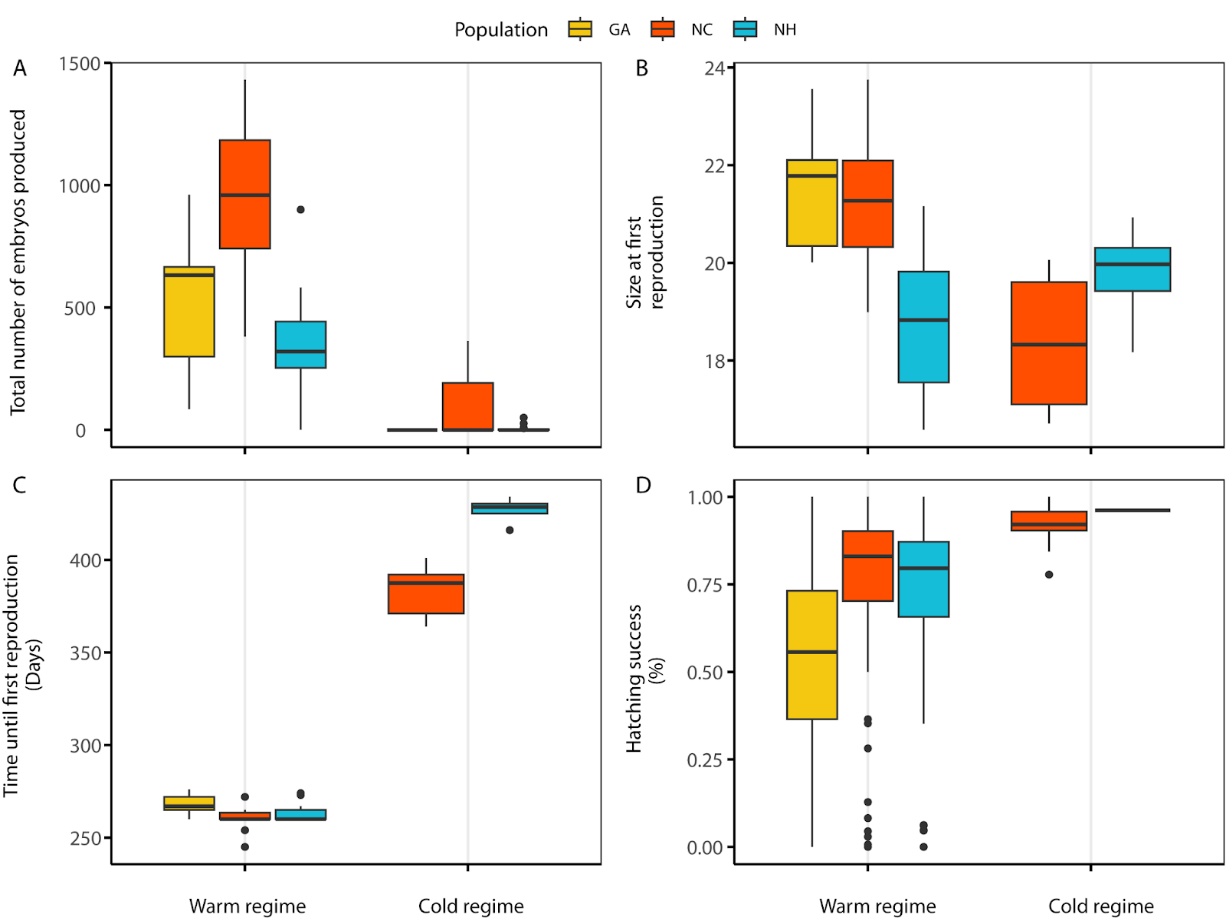

### Figure S2:

Reproductive trait data incorporating Georgia population (gold). Graphics for each panel depict measurements for each population as described for Figure 4. Note that in the cold regime, no females from the Georgia population reproduced during the experiment.
